# Discrete somatotopic representation and neural response properties in hummingbird and zebra finch forebrain nuclei

**DOI:** 10.1101/2023.10.30.564821

**Authors:** Andrea H. Gaede, Pei-Hsuan Wu, Duncan B. Leitch

## Abstract

Somatosensation allows animals to perceive the external world through touch, providing critical information about physical contact, temperature, pain, and body position. Somatosensory pathways, particularly those related to the rodent vibrissae, have been well-studied in mammals, illuminating principles of cortical organization and sensory processing ^1,2^. However, comparative studies across diverse vertebrate species are imperative to understand how somatosensory systems are shaped by evolutionary pressures and specialized ecological needs.

Birds provide an excellent model for studying the evolution of somatosensation, as they exhibit remarkable diversity in body plans, sensory capabilities, and behavior. Prior work in pigeons^3-6^, parrots^7^, and finches^8^ have identified general tactile-responsive regions within the avian telencephalon. Yet how somatosensory maps and response properties vary across key avian groups remains unclear. Here, we aimed to elucidate somatotopic organization and neural coding in the telencephalon of Anna’s hummingbirds (*Calypte anna*) and zebra finches (*Taeniopygia guttata)*.

Using *in vivo* extracellular electrophysiological techniques, we recorded single and multi-unit activity in telencephalic regions of anesthetized hummingbirds and finches. We stimulated the beak, face, trunk, wings, and hindlimbs with controlled tactile stimuli and mapped somatosensory receptive fields. We found distinct representations of body regions distributed across multiple somatosensory zones, with surprising differences in relative areas devoted to key body surfaces, potentially as related to behavioral significance.

**Highlights:**

□ Somatosensation provides birds with critical information for behaviors necessary to survival including foraging and flight.
□ *In vivo* extracellular physiological recordings were used to monitor tactile responses in contralateral forebrain nuclei corresponding to the feather deflection in hummingbirds and finches including to air puff stimuli.
□ Both hummingbirds and finches show distinct separation of body and head receptive field representation in different nuclei.
□ A continuous somatotopic arrangement can be found in both the rostral Wulst (corresponding to the wings and body) and in nucleus basorostralis (corresponding to the head and beak), with particularly enlarged representations of the wing leading edge and the foot.

## Results

Hummingbirds are specialist nectarivores with unique demands on tactile senses for precision hovering and feeding **(Fig. 1A)**. Their sensory systems are tuned for fast and nuanced control of flight and specialized tongue and bill movements^9-13^. In contrast, songbirds like finches use a flap-bounding flight style and employ their bill primarily for foraging and manipulation **(Fig. 1B)**.

**Figure 1:**
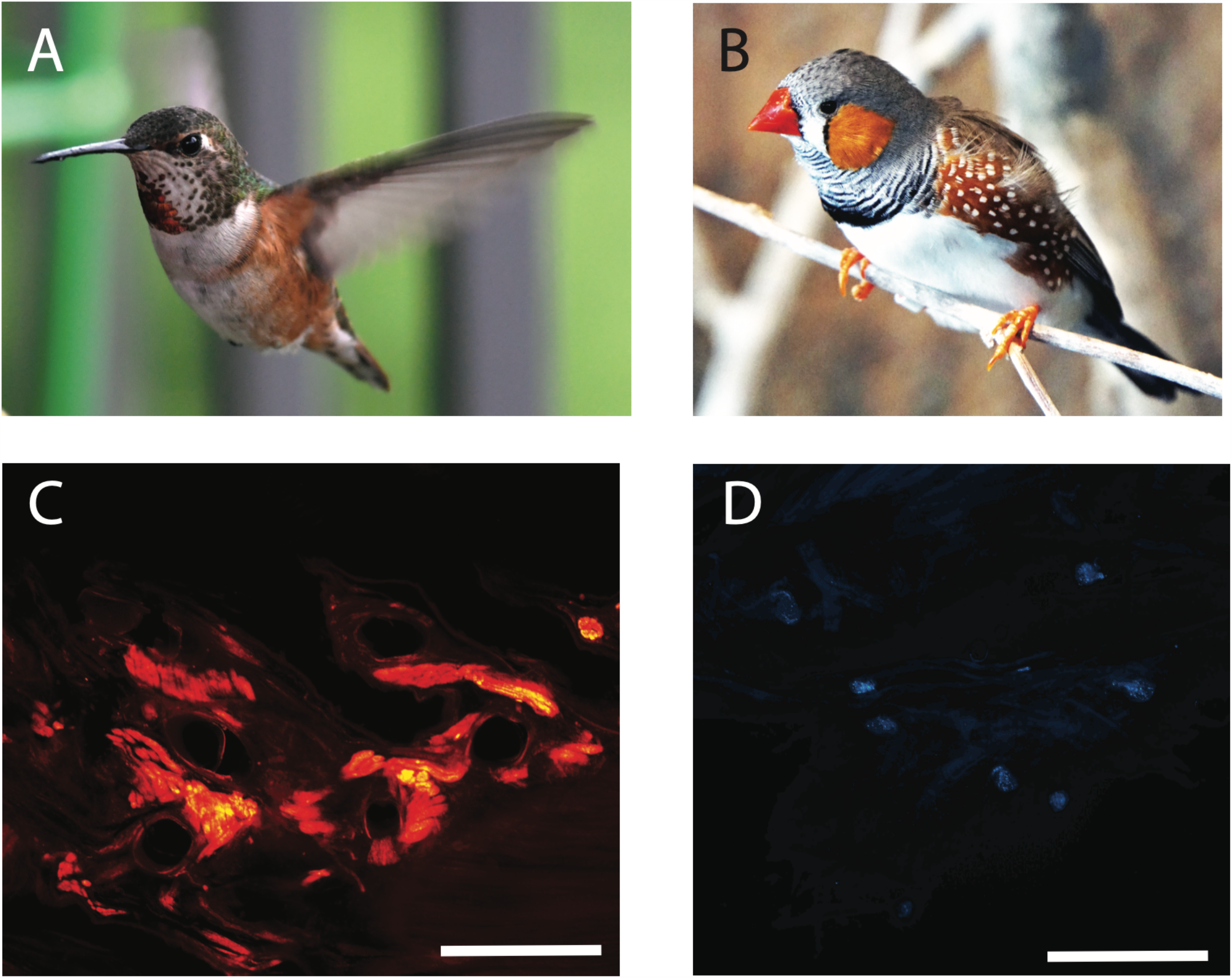
Comparative anatomy of avian tactile systems. **(A)** Hummingbirds have sensory adaptations (e.g., optic flow analysis) facilitating hovering flight. Photo by authors. (B) Zebra finches employ intermittent flight while foraging. Photo used under a Creative Commons license. (C) DiI labeling, applied to the hummingbird radial nerve, shows the arcade of myelinated fibers at the feather/dermis juncture. Scale = 1 mm. (D). Putative mechanosensory Herbst corpuscles labeled with AM1-43 in a hummingbird wing, following mechanical stimulation. Scale = 1 mm.

### Peripheral mechanoreceptor structures are associated with feathers of the wing and glabrous surfaces

We hypothesized that feathers of the wing, including those associated with filoplume and leading edge feathers would show particular innervation by mechanosensory end organs. Herbst corpuscles (avian specific homologs of Pacinian lamellated corpuscles of mammals and other vertebrates) have been associated with the non-feathered skin of the beak in ducks^14^, kiwis^15^, and shore-foraging birds^16,17^ that appear to localize prey using vibration detection via specialized sensory pits at the rostral margins of the beak. Along feather-covered surfaces, Herbst corpuscles are associated with the follicles of facial bristle feathers^18^, conspicuous whisker-like feathers that appear from margins of the mouth and above the eyes in diverse lineages of birds including kiwis^19^ and owls^20^.

We performed anterograde tracing in the wings of 4 hummingbirds, applying the lipophilic dye DiI (1, 1’-dioctadecyl-3,3,3’3’-tetramethylindocarbocyanine) to the proximal ends of cut nerves of the brachial plexus including the radial nerve. At the base of the leading edge of the wing, the follicles of the primary and filoplume feathers are well innervated by a ring-like “arcade” network^21^ of large myelinated fibers that extend into the dermis (**Fig. 1C**). Finer unmyelinated free nerve endings extend to more superficial layers of the epidermis.

We next analyzed how mechanoreceptors are distributed across the leading edge of the wing surface. *In vivo* subcutaneous injections of fluorescent AM1-43, followed by anterior to posterior tactile stimulation of wing (mimicking air flow patterns during flight) were used to visualize sensory neurons and their location across areas of the leading edge of the wing. Distinct swellings associated with Herbst corpuscles were sparse but enriched in proximity to the feather follicle complexes (**Fig. 1D**). This conformation suggests that these homologs of vibration sensitive Pacinian corpuscles are potentially well-positioned to detect mechanical deflection associated with diverse flight behaviors, including stall detection.

### Central organization of the rostral Wulst

Visually responsive areas of the hyperpallium (Wulst) receive input via the thalamofugal system, whereas somatosensory Wulst is the target of ascending fibers from a dorsal thalamic nucleus (nucleus dorsalis intermedius ventralis anterior, DIVA)^22,23^. DIVA itself is the target of the dorsal column nuclei^24^.

In both hummingbirds and finches, we identified single and multi-unit receptive fields corresponding to mechanical stimulation of the contralateral non-facial body surface. The somatosensory Wulst includes the most superficial lamina of Wulst (hyperpallium; H, located medially at rostral levels), then ventrolaterally the narrow somatosensory thalamo-recipient lamina called the intercalated hyperpallium (IH), and finally the dorsal mesopallium (MD) more laterally^25,26^ **(Fig. 2A-B)**. Tactile responses, extending from the upper back/nape of the neck to the tail feather and hindlimb were marked on photographs of the body, following repeated manual stimulation.

**Figure 2:**
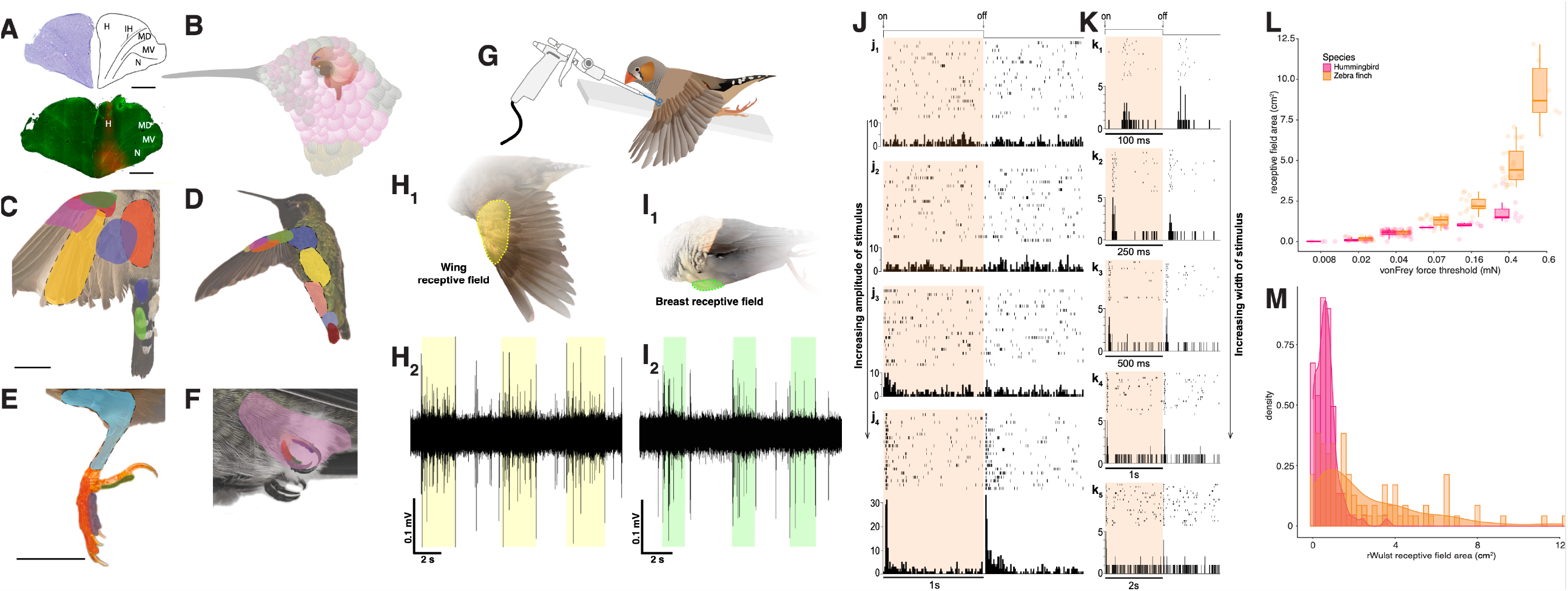
Central organization and response properties of the somatosensory Wulst. (**A**) Photomicrographs of coronal sections through the hyperpallium. Top: Nissl-stained coronal section (top, left) and anatomical borders (top, right) in the rostral telencephalon of the zebra finch. H = hyperpallium, IH = intercalated hyperpallium, MD = dorsal mesopallium, MV = ventral mesopallium, N = nidopallium. Scale bar = 1 mm. Bottom: Photomicrograph showing recording site (red) in the rostral Wulst of the finch. Fluorescent Nissl stain (green) used to identify anatomical borders. H = hyperpallium, MD = dorsal mesopallium, MV = ventral mesopallium, N = nidopallium. Scale bar = 1 mm. (**B**) Illustration showing the orientation of the brain and rostral (somatosensory) Wulst in Anna’s hummingbird. (**C**) Example tactile receptive field borders distributed over the wing in the finch and (**D**) the hummingbird. Scale bar is 1 cm. (**E**) Example tactile receptive field borders distributed over the hindlimb in finch and F) the hummingbird. Scale is 1 cm. (**G**) Illustration of experimental prep for delivering naturalistic airflow. Small puffs of compressed air (1-2s) were delivered to identified receptive fields while simultaneously recording single units in the rostral Wulst. (**H**) Wing receptive field comprised of coverts and alula (**H**_**1**_) defined by yellow shading. Raw neural trace (**H**_**2**_) from rostral Wulst corresponding to cell with receptive field shown in **H**_**1**_. Yellow highlighted region of trace indicates stimulus ‘on’ period, i.e., air puff is applied to the receptive field during the yellow period. Cell activity is increased during the air puff stimulus. (**I**) Finch breast receptive field defined by green shading (**I**_**1**_). Raw neural trace from rostral Wulst neuron (**I**_**2**_) corresponding to receptive field shown in **I**_**1**_. Green highlighted region of trace indicates stimulus ‘on’ period, i.e., air puff is applied to the receptive field during the green period. Cell activity is increased during the air puff stimulus. (**J**) Raster plots and peri-stimulus time histograms (PSTH) for finch rostral Wulst neuron in response to computer-controlled piezo-electric mechanical stimulus. Amplitude of stimulus increases from 0.5 mm (**j**_**1**_) to 2 mm (**j**_**4**_). Orange panels indicate stimulus ‘on’ period. (**K**) Raster plots and PSTH summarizing finch rostral Wulst neuron responses to a computer-controlled piezo-electric mechanical stimulus with increasing stimulus ‘on’ period. Stimulus period increases from 100 ms (**k**_**1**_) to 2 s (**k**_**5**_). Orange panels indicate stimulus ‘on’ period. (**L**) Receptive field surface area for rostral Wulst cells decrease as sensitivity to force increases. Quartile boxplots and raw data display receptive field size for each hummingbird or zebra finch rostral Wulst neuron at the minimum force detected by that cell. Receptive field surface area generally increased faster for finches than for hummingbirds. Magenta = hummingbird, orange = zebra finch. (**M**) Histogram with kernel density plot overlay showing the prevalence of receptive field areas for each species. Hummingbird receptive field sizes are clustered at smaller areas than finches, which are observed at a broad range of sizes. Magenta = hummingbird, orange = zebra finch.

With increasing depth of the electrode along the dorsoventral axis, we found a gradual shift in body tactile representation. For example, in a single finch Wulst recording track, receptive fields gradually progressed from the upper back/nape (3100 *μ*m from surface), to the shoulder and wing (3300 *μ*m), to mid back (3500 *μ*m), and finally to the tail feathers (3700 *μ*m) **(Fig. 2C)**. Similarly, representative hummingbird receptive fields from a single track are shown in **Fig. 2D**. Across most of its rostrocaudal axis, the Wulst was approximately 600 *μ*m in height. In the hummingbird, the rostral Wulst was approximately 400 *μ*m in height. This general conformation of the dorsoventral axis corresponding broadly to the anterior-posterior axis of the body was also found in representations of the hindlimb, with receptive fields corresponding to the foot feathers/ankle gradually progressing to representations of the ventral surface of the foot with increasing depth.

Across the rostral Wulst’s long axis, the non-facial body was also represented modularly, with broad areas corresponding to neck, breast, wing, foot surface, and tail. Similarly, in the hummingbird rostral Wulst, these same areas were well-represented. Notably, in both species, there were many distinct receptive fields corresponding to the glabrous skin surface of the hindlimb **(Fig. 2E, F)**. In hummingbirds, receptive fields related to the foot accounted for 17.8% of Wulst responses, whereas in finches, foot-related fields accounted for 10.1% of responses.

### Responses to naturalistic airborne stimuli

We next investigated whether multiunit responses in the finch Wulst could be elicited from semi-naturalistic air puff stimuli to mimic possible aerial contact associated with wind gusts or flight. After identifying the general location of a receptive field via manual contact, we positioned an air puff stimulus approximately 2 cm away from the center (**Fig. 2G**). These body areas included feathers of the chest and the wing, including the leading edge and coverts.

Indeed, robust bursts of spiking were elicited as each air puff was released upon the feathered surface (**Fig. 2H-I**). Furthermore, changing the angle of the air stimulus appeared to modulate the latency and spike density of the response, possibly indicating a role for directional encoding of gusts or feather position. Systematic measurements of air puff strength and direction, with respect to field position across the body, are necessary to draw stronger conclusions about the tactile information provided by aerial stimuli as processed in the somatosensory Wulst.

### Response properties of the rostral Wulst

For single unit-recordings, once the general receptive field location had been determined using manual tactile stimulation, a piezoelectric stimulator was positioned within the center of the receptive field and single-unit activity was recorded (**Fig. 2J, K**). Spike sorting revealed that the majority of responses recorded in both hummingbird (94/123) and finch (187/243) rostral Wulst continued to respond to the ongoing indentation of the receptive field (i.e., slow adaptation) for all stimuli lengths (100 ms to 2 s). In response to increasing indentation from the piezo stimulus (from 0.5 mm to 2 mm), neurons responded with increasing firing frequency.

Along with measurement of excitatory response pattern, we assessed body tactile sensitivity using calibrated von Frey filaments. In hummingbird Wulst, mechanical thresholds of activation ranged from 8 mN (the lowest calibrated amplitude) to 400 mN, whereas in the finch these measurements varied from 20 to 600 mN (**Fig. 2I**). In both species, the lowest threshold responses were observed on the surface of the hindlimb and the elongated tail feathers, and the highest amplitude thresholds corresponded to areas of the back and scapula. In a hummingbird case, we also found a receptive field on the leading edge of the wing that responded to 8 mN deflection.

We measured the surface area of receptive fields on photographs of the extended body surface. For hummingbirds, the smallest receptive fields were found on the leading edge of the wing (0.008 cm^2^) and foot surface (0.01 to 0.035 cm^2^) and the largest (0.58 to 3.59, 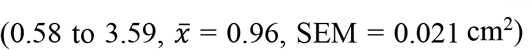, SEM = 0.021 cm^2^) were found on the back. Finches demonstrated a similar pattern of the smallest receptive fields corresponding to the foot (0.043 to 0.16 cm^2^) and largest fields related to the back (2.18 to 12.17, 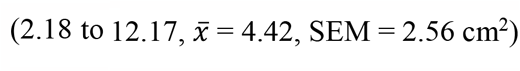, SEM = 2.56 cm^2^).

Examining this relationship between area and sensitivity, we found that receptive field surface area and von Frey threshold were significantly correlated. With increasing von Frey force threshold, receptive field size generally increased faster for finches than for hummingbirds. (**Fig. 2L**). Hummingbird receptive field areas were primarily clustered in smaller sizes (under 2 cm^2^) compared to the wider range of areas observed in zebra finches (**Fig. 2M**).

### Central organization of nucleus basorostralis

As the direct target of the principal sensory nucleus of the trigeminal nerve via the quintofrontal tract, the nucleus basorostralis is a multimodal nucleus in the avian ventral anterior telencephalon (**Fig. 3. A-C**)^5,24^. Given the variation of body regions represented in somatotopic responses, from the beak and bill alone in pigeons^5^ and finches^8^ to the entire body surface in barn owls^27^ and budgerigars^7^, we wondered to what extent the hummingbird basorostralis followed either pattern. We hypothesized that hummingbirds may show expanded representation or enhanced sensitivity in trigeminal projections encoding beak inputs. Secondarily, we investigated whether auditory responses were interdigitated within somatosensory representations, as in the case of the budgerigar^7^.

**Figure 3:**
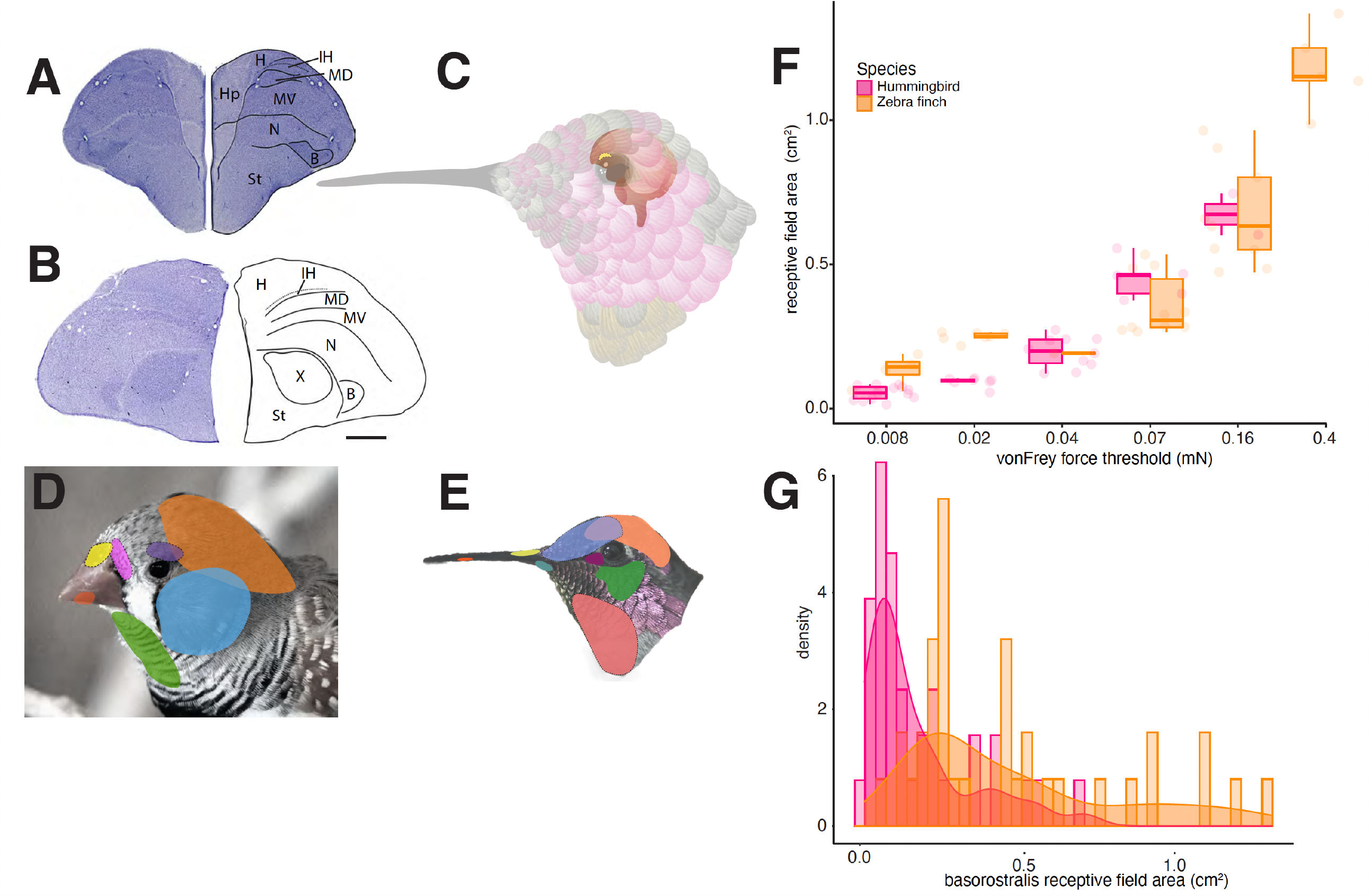
Central organization and response properties of nucleus basorostralis. Photomicrographs of Nissl-stained coronal section and anatomical borders in the rostral telencephalon of the hummingbird (**A**) and zebra finch (**B**). H = hyperpallium, IH = intercalated hyperpallium, MD = dorsal mesopallium; MV = ventral mesopallium, N = nidopallium, B = basorostralis, St = striatum, Hp = hippocampus, X= area X. Scale bar = 1 mm. (**C**) Illustration showing the orientation of the brain and nucleus basorostralis (B) in Anna’s hummingbird. (**D**) Example tactile receptive fields distributed over the head and beak in finches and in (**E**) hummingbirds. (**F**) Receptive field size for Bas cells decrease as sensitivity to force increases. Quartile boxplots and raw data display receptive field size for each hummingbird or zebra finch Bas neuron at the minimum force detected by that cell. Magenta = hummingbird, orange = zebra finch. (**G**) Histogram with kernel density plot overlay showing the prevalence of receptive field areas for each species. Magenta = hummingbird, orange = zebra finch.

In both the hummingbird and finch, we found representations to be dominated by areas innervated primarily by the trigeminal system. These areas included the caudal portion of the head, areas adjacent to the eyes, as well as the neck (**Fig. 3 D, E**) Furthermore, at more anterior areas of basorostralis, we identified receptive fields corresponding to the beak, with these progressing from proximal to distal areas of the beak as the electrode was placed gradually more rostral. We did not readily identify any areas responding to auditory stimulation. Furthermore, all receptive fields within basorostralis appeared to correspond to the surface of the head, neck, and beak, with no distinct responses to tactile stimulation of the post-cranial body surface.

### Response properties of nucleus basorostralis

Similar to our examinations of rostral Wulst, we employed *in vivo* extracellular electrophysiology to assess both receptive field surface area and tactile force thresholds. When recording from the finch basorostralis and employing a sustained square wave tactile stimulus (indentation movement of 1 mm) from a piezoelectric device, 74% of neuronal responses (14/19) appeared to be rapidly adapting or phasic, with an excitatory response to onset and offset of the 500 ms stimulus. In contrast, 26% of responses (5/19) appeared to slowly-adapted or tonic, with ongoing spiking across the duration of the stimulus presentation.

Using von Frey filaments, we assessed the minimal amount of mechanical force needed to elicit spiking. In hummingbirds, we found that fields spanning the beak, cheek, chin, and directly adjacent to the eye had thresholds corresponding to the finest filaments (8 mN) whereas areas corresponding to the posterior surface of the head required greater force (70 to 160 mN). Similarly, recording in finch basorostralis, we found that 8 mN elicited activity across beak, chin, and throat surfaces, and the crown area of the dorsal surface of the head required the greatest measured indentation (400 mN) (**Fig. 3F**).

In both finches and hummingbirds, basorostralis receptive fields of the smallest surface areas were recorded across the beak and chin. For example, our smallest receptive field (0.015 cm^2^) was found on the hummingbird beak. The largest receptive fields were found on the back surface of head in both species, along with large receptive fields related to the crown of the head in the finch (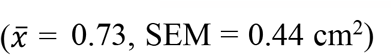, SEM = 0.44 cm^2^) (**Fig. 3G**). Examining this relationship between area and sensitivity, we found that receptive field surface area and von Frey threshold were significantly correlated (**Fig. 3F**).

## Discussion

We investigated forebrain representation of tactile body surfaces in hummingbirds and finches, with particular focus on somatotopic organization and response properties, employing *in vivo* electrophysiological techniques. Given the diversity of somatotopic arrangements that have been identified among classical models of mechanotransduction (e.g., rodents)^28,29^, we were interested in exploring whether similar somatosensory maps are present within birds – the most diverse group of land vertebrates with more than 10,000 extant species^30^.

### Avian somatosensory system representation

One key finding of our study is the identification of clearly defined tactile receptive fields within the avian forebrain. Building from the seminal work of Wild and colleagues who recorded multiunit responses from Wulst and basorostralis in songbirds^8^, owls^27,31^, ducks^32^, among others (**Fig. 4A**), we employed single unit analyses, observing robust neuronal responses in specific regions during mechanical stimulation, indicating the presence of dedicated tactile processing circuits. These nuclei were characterized by receptive fields that were spatially and anatomically segregated, suggesting a topographic representation of tactile information in the avian forebrain (**Fig. 4B-C**). Specifically, we found that areas corresponding to the head, neck, and beak were robustly represented within nucleus basorostralis. In both hummingbird (**Supplemental video 1)** and finch (**Supplemental video 2)**, these representations followed a generalized pattern, with caudal areas of the head found in the posterior margins of basorostralis with gradual progression to the beak representation, which occupied the anterior half of basorostralis. Receptive fields corresponding to dorsal regions (e.g., the maxillary beak) were typically found superior to ventral regions (e.g., the mandibular beak). Our recording preparation, in which the anesthetized bird was held stationary with its beak closed precluded detailed mapping of intraoral regions or the tongue, an area that deserves further investigation.

**Figure 4:**
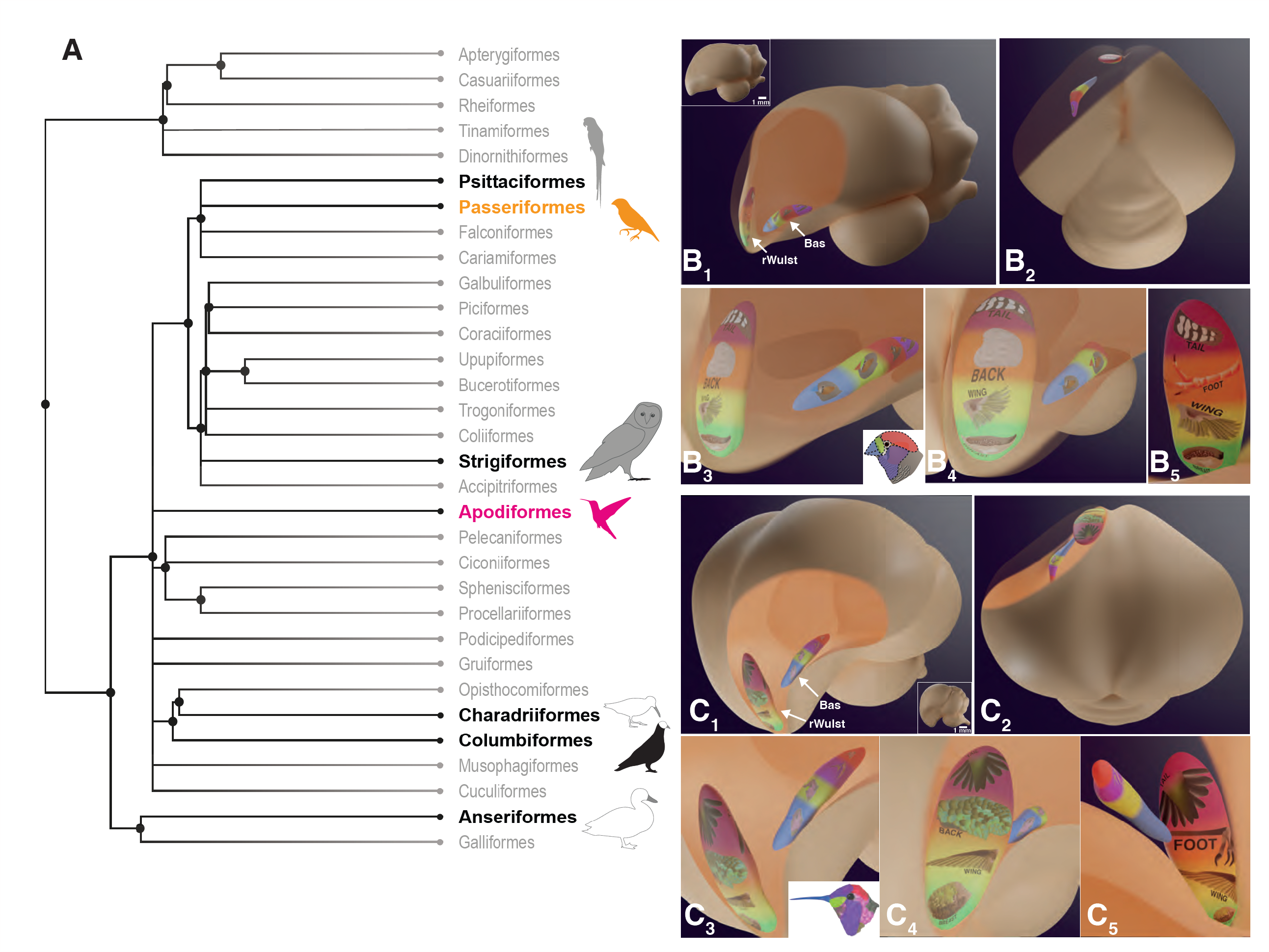
Comparative view of the avian somatosensory system representation. (**A**) Illustration of phylogenetic relationships between avian orders. Black text indicates groups for which somatosensory data from the rostral Wulst or nucleus basorostralis has been reported. Animal illustrations provide a general indication of which body parts are represented in each nucleus (gray fill = basorostralis, black fill = rostral Wulst); e.g., budgerigars (Psittaciformes)^7^ have representations of the body and beak in basorostralis, barn owls (Strigiformes) have a representation of the claw in the rostral Wulst^31^,^44^ and representation of the body and beak in basorostralis, while pigeons (Columbiformes) have a beakless representation in the rostral Wulst^4^ and the beak is represented in basorostralis^6,45^. Groups used in the present study are highlighted in orange (zebra finches) and magenta (hummingbirds). (**B**) 3D illustrations of the zebra finch brain with interior views showing relative size and position of rostral Wulst (rWulst) and basorostralis (Bas) in the left hemisphere. Color indicates region of body represented. For rWulst: magenta = tail, anterior orange = back, posterior orange = foot, yellow = wing, and green = breast. For Bas: blue = beak, yellow-green = near beak, red = crown, purple = cheek, chin, throat. (**B**_**1**_) Sagittal view of the finch brain, rostral pole to the left, caudal pole to the right. Inset shows a sagittal view of the brain with no opacity so that gross anatomy can be visualized. (**B**_**2**_) Dorsal view of finch brain. (**B**_**3**_) Magnified view of the rostral telencephalon illustrating body representation in the rWulst and beak and head representation in Bas. Note that both color and drawings of the body/head region indicate the area represented in each nucleus. The inset shows how regions of the head are represented in Bas. (**B**_**4**_) Magnified view of rostral telencephalon; more anterior view than B_3_. (**B**_**5**_) Caudal view of rWulst. Image acquired from inside the brain looking anteriorly. Note, from this angle, the foot representation is visible. (**C**) 3D illustrations of the hummingbird brain with interior views showing relative size and position of rWulst and Bas in the left hemisphere. Color indicates region of body represented. For rWulst: magenta = tail, anterior orange = back, posterior orange = foot, yellow = wing, and green = breast. For Bas: blue = beak, yellow-green = near beak, chin, red = crown, purple = cheek, throat, near eye. (**C**_**1**_) ¾ view of the hummingbird brain with relative position of rWulst and Bas visible. Inset shows a sagittal view of the hummingbird brain with no opacity so that gross anatomy can be visualized. Rostral pole is to the left, caudal pole to the right. (**C**_**2**_) Dorsal view of hummingbird brain. Note that the brain model has been pitched backwards to give a clear view of rWulst and Bas. (**C**_**3**_) Magnified view of the rostral telencephalon illustrating the body representation in rWulst and beak and head representation in Bas. Note that both color and drawings of the body/head region indicate the area represented in each nucleus. The inset shows how regions of the head are represented in Bas. (**C**_**4**_) Magnified view of the rostral telencephalon; more anterior view than C_3_. (**C**_**5**_) Caudal view of rWulst. Image acquired from inside the brain looking anteriorly. Note, from this angle, the foot representation is visible.

Similar to the arrangement noted in pigeons^3,4^, the rostral Wulst of both hummingbirds and finches encompassed areas exclusively corresponding to the post-cranial body. Along its rostrocaudal axis, the Wulst featured receptive fields related to the breast, wing, back, and tail feathers, respectively. Following this longest axis of the Wulst in both species, we also encountered relatively large areas dedicated to tactile responses from the glabrous surface of the foot and adjacent feathers of the foot. This was positioned directly adjacent to areas corresponding to the back and tail feathers. This distinct Wulst organization of the surface of the foot and associated regions is reminiscent of the prominent claw representation found in the somatosensory Wulst of the barn owl^31^, a species which notably employs its talons and foot surface in securing prey. Whereas hummingbirds and finches do not immobilize prey using their feet, these sensory surfaces are used in a variety of important tactile tasks. These include spending significant amounts of time perched in branches, with their feet well-adapted for gripping surfaces, nest building, preening, and scratching. In addition, finches routinely use their feet while feeding, holding securing vegetation or seeds for cracking with the beak. Further detailed examination of the proportional volume of the body representation within Wulst could shed light on whether this notable hindlimb representation shows similarities to the enhanced representation of discrete body surfaces, as observed primarily in mammalian primary somatosensory cortex. This phenomenon of “cortical magnification” – the preferential allocation of the cortical real estate to behaviorally significant sensory receptors such as the surfaces of the hands and lips in primates reflected in primary somatosensory cortex^33^ – is thought to contribute to increased sensory resolution for important areas of the body periphery. As this appears to be the case in the barn owl (to the extent that the body is no longer represented within the somatosensory Wulst), further physiological research employing falcons, hawks, and parrots^34^, all birds that have been shown to use their feet for particularly complex manipulations, could shed light on more specific granular representation of the foot surface within the Wulst. Physiological and anatomical examinations of the nucleus basorostralis in the dunlin (an avian probe-feeding specialist)^35^ suggest that preferential expansion in representation of important sensory surfaces (e.g., the bill tip) might potentially be found among other birds.

### Receptive field properties

Avian mechanotransduction differs from mammalian systems in several fundamental ways^36^. Forebrain somatosensation is processed in in the hyperpallium and discrete nidopallial nuclei, ventral to the superficial pallium, whereas primary somatosensory cortex (S1) is distributed across the laminated neocortex. At the periphery, mammalians rely upon diverse classes of mechanoreceptor end organs (e.g., Meissner corpuscles, Merkel complexes) and tactile organs associated with the follicle complex of specialized vibrissa. Although birds also have some classes of homologous mechanoreceptor end organs (e.g., Herbst and Grandry corpuscles), the majority of their bodies are covered by distinct populations of diverse types of feathers. Therefore, we wondered how basic receptive field properties including area and mechanical threshold might compare to classically studied somatosensory models, as measured from the relays within the avian forebrain.

In general terms, smaller receptive fields are associated with higher sensitivity. Smaller receptive fields allow for greater spatial resolution, meaning the ability to discern fine details about the stimulus. For example, human fingertips and the lips have minute receptive fields and a high density of touch receptors, providing peripheral adaptations for high sensitivity and precision in tactile discrimination^37^.

On the other hand, larger receptive fields are typically associated with lower sensitivity. Areas of the body including the back or upper hindlimbs have larger receptive fields and fewer mechanoreceptors, which results in lower tactile sensitivity and spatial resolution. This inverse relationship between receptive field surface area and absolute mechanical thresholds of sensitivity has been explored in other vertebrates, including in close avian relatives, the archosaurs^38^. Given the prominent representation of the claw in both the owl basorostralis^31^ and the foot in the rostral Wulst in hummingbirds and finches, combined with recent insight into the diversity of foot usage among diverse bird lineages^34^, further investigation of tactile specialization of the avian hindlimb and its representation within the telencephalon could yield insight into general principles of sensory adaptation.

It is worth noting that our study focused primarily on the avian forebrain, and further investigations are warranted to explore tactile processing in other avian brain regions, including the brainstem and diencephalon. Prior investigations using extracellular radial nerve recordings, corresponding to mechanical and air puff stimuli in chickens, suggest that feathers can respond robustly to airflow stimuli of varying directions^39^. Further neurophysiological recordings along the somatosensory neuroaxis might potentially reveal segregation in processing differing forms of somatosensory input (e.g., pain vs. mechanical deflection), as has been indicated within trigeminal brainstem relays in mammalians^40^. Additionally, future studies could employ more advanced techniques, such as *in vivo* functional imaging, to gain a more comprehensive understanding of the neural dynamics and circuitry underlying avian tactile processing. Indeed, molecular techniques have made progress in identifying ion channel contributions to tactile and thermal sensation, using the unique distribution of sensory end organs within the duck beak^41^. Similar approaches applied comparatively across avian taxa^42^ may further illuminate diversity in sensory ion channel function, as well as more broadly the evolutionary interplay between sensory structures, neural processing, and behavioral adaptation.

In conclusion, our investigation provides novel insights into tactile receptive field organization and response properties in the avian forebrain, using hummingbirds and finches, which both vary in their tactile behavior and ecology. By delineating architectural principles and functional properties of avian tactile processing in the hyperpallium and nucleus basorostralis, we contribute to the broader field of comparative neurobiology and enhance our understanding of sensory systems. Future research in this area has the potential to unveil further intricacies of avian tactile perception and shed light on the remarkable sensory capabilities of birds.

## Methods

Electrophysiological recordings and histological images were acquired from six adult male Anna’s hummingbirds (*Calypte anna*) and 20 adult male zebra finches (*Taeniopygia guttata*). All animal procedures were approved by the University of British Columbia Animal Care Committee in accordance with the guidelines set out by the Canadian Council on Animal Care or conducted according to Home Office guidelines under the UK Animals in Scientific Procedures Act 1986.

### *In vivo* labeling

Feather-covered hummingbird skin samples were removed post-mortem from paraformaldehyde (PFA)-fixed tissues. Small crystals of DiI (1, 1’-dioctadecyl-3,3,3’3’-tetramethylindocarbocyanine, Invitrogen, Carlsbad, CA) were applied via insect pins to distal branches of spinal nerves innervating the wings, as dissected from the brachial plexus. These samples were embedded in 2% agarose, immersed in 4% PFA, and stored in darkness at 27°C for at least 4 weeks.

In other hummingbirds, AM1-43 (Biotium, Fremont, CA) was diluted in sterile PBS and injected subcutaneously near the convergence of the wing and back. At 22 hours post-injection, the wing feathers were manually stimulated via periodic brushing for one hour. All specimens were sectioned sagittally at 80 uM thickness on a sliding microtome (Leica, Concord, Ontario, Canada) and imaged on a Zeiss AxioImager M2 (Zeiss, Jena, Germany).

### Electrophysiological measurements

Stereotaxic surgery was performed using an adapted frame (David Kopf) that secured the head of the hummingbird or zebra finch. Coordinates were derived from Nissl-stained sections and from neuroanatomical measurements in zebra finch basorostralis^8^ and rostral Wulst^4^. Birds were anesthetized with an intramuscular injection of ketamine/xylazine (65 mg/kg ketamine, 16.7 mg/kg xylazine, i.m.) and supplemental injections to maintain a surgical plane were given as needed. Digital photographs were made of the body surface of each bird to serve as reference for distinguishing boundaries of receptive fields. Initially, the head was angled downward at 45° to the horizontal plane to access the contralateral anterior telencephalon via small craniotomy (approximately 1.5 mm x 1.5 mm), and the dura mater was removed.

Single and multi-unit neuronal recordings were performed using glass microelectrodes (5 μm diameter tip), filled with 2M NaCl, or via 20 MOhm tungsten electrodes (FHC, Bowdoin, ME). A silver wire clipped to the skin adjacent to the incision served as a reference electrode. The recording electrode was attached to a micro manipulator (Sutter, Novato, CA), oriented perpendicularly to the surface of the telencephalon, and advanced while listening to audio output of neural activity in response to gentle mechanical stimuli.

The locations of recording sites were confirmed via dextran microinjection (Texas Red 3000 MW or micro-Emerald 3000 MW, MilliporeSigma, Ontario, Canada). At the end of each experiment, the bird was euthanized via overdose of ketamine/xylazine and transcardially perfused with 0.9% saline followed by 4% PFA. Brains were dissected, cryo-protected in 30% sucrose for at least 3 days, and sectioned (40 μm) coronally. Tissue sections were stained for Nissl substance (NeuroTrace 500/525, ThermoFisher; thionin, MilliporeSigma) to visualize forebrain architecture, and the location of the dextran injection was verified using a Zeiss AxioImager M2 (Zeiss Research Microscopy, Jena, Germany).

### Tactile stimuli

We recorded multi-unit neural responses in response to manual gentle brushing and tapping with wooden probes to investigate the boundaries of tactile receptive fields. We used calibrated plastic filaments (von Frey hairs, Stoelting, IL) to record the mechanical threshold required to elicit neural responses in the telencephalon. These receptive fields were manually outlined on digital photographs of the body surface of each bird.

Upon recording from a well isolated unit, a square wave stimulator (Powerlab, AD Instruments, Sydney, Australia) was used drive a mechanical probe attached to a E-625 Piezo Servo Controller (Physik Instrument LP, Auburn, MA) which was position in the center of the tactile receptive field. The amplitude and length of the square pulse were independently adjusted.

In a second cohort of finches, we also used airflow stimuli, using a custom-built compressor system. After identifying and mapping each receptive field as described above, the airstream was positioned perpendicular to the receptive field, approximately 2 cm from the surface, and short (1-2s) air bursts were applied. Subsequent trials applied air flow at a range of angles (e.g., simulating forward flight or a wind gust from behind).

### Data collection and analysis

Neural activity was amplified (x10,000) (A-M Systems, Sequim, WA), bandpass filtered (0.1 to 3 kHz), and acquired at 20 kHz using a CED micro1401-3 (Cambridge, UK). We processed raw neural recordings offline using the spike sorting algorithm in Spike2 (Cambridge Electronic Design, Cambridge, UK) to extract single units. This program enabled us to set trigger thresholds and dimensions of a sliding window encompassing the full spike amplitude to identify individual spikes. These spikes were then matched to full wave templates for classification. We grouped similar templates post-hoc using principal component analysis and visual inspection of overlaid spikes coded by template. Spike sorted data were used to produce raster plots and peri-stimulus time histograms to visualize neural responses to tactile stimuli.

### Receptive field quantification

Prior to physiological recordings, we collected digital photographs of the body surface of each hummingbird or finch with a surgical scale bar. Post-surgery, we outlined the borders of drawn receptive fields, calibrated to the scale bar using Fiji software, to measure surface area^43^.

